# PhaB-independent poly(3-hydroxybutyrate) production in the thermophilic hydrogen-oxidizing bacterium *Hydrogenophilus thermoluteolus*

**DOI:** 10.64898/2026.05.08.723683

**Authors:** Kotaro Yoda, Masafumi Kameya, Hiroyuki Arai

**Affiliations:** Social Cooperation Program for Next-generation Biomanufacturing, The University of Tokyo, Yayoi 1-1-1, Bunkyo-ku, Tokyo 113-8657, Japan; Department of Biotechnology, The University of Tokyo, Yayoi 1-1-1, Bunkyo-ku, Tokyo 113-8657, Japan; Collaborative Research Institute for Innovative Microbiology, The University of Tokyo, Yayoi 1-1-1, Bunkyo-ku, Tokyo 113-8657, Japan

**Author notes:** Corresponding author. E-mail address (H. Arai).

**Keywords:** β-oxidation, hydrogen-oxidizing bacterium, metabolic engineering, poly(3-hydroxybutyrate), *Hydrogenophilus thermoluteolus*

## Abstract

*Hydrogenophilus thermoluteolus* TH-1 is a thermophilic hydrogen-oxidizing bacterium capable of producing poly(3-hydroxybutyrate) (PHB) from CO_2_. To redirect carbon flux for producing other useful biomaterials, we disrupted the acetoacetyl-CoA reductase genes (*phaB1* and *phaB2*), which are central to the primary PHB synthesis pathway. Unexpectedly, the resulting Δ*phaB1B2* mutant still accumulated PHB under autotrophic conditions, reaching approximately 25-35 % of the wild-type level. Furthermore, PHB accumulation in the mutant was significantly restored when fatty acids (butyrate and oleate) were used as carbon sources, whereas acetate and malate resulted in reduced accumulation. These results suggest the existence of a PhaB-independent PHB synthesis pathway. We propose that intermediates from the β-oxidation of fatty acids are converted to (*R*)-3-hydroxybutyryl-CoA, bypassing the disrupted PhaB enzymes. Additionally, the basal PHB production from non-fatty acid sources implies the involvement of a reverse β-oxidation pathway. This study highlights the metabolic versatility of strain TH-1 for future metabolic engineering.

## INTRODUCTION

The conversion of carbon dioxide (CO_2_) into biomaterials by microorganisms is an attractive strategy for the utilization of CO_2_, a well-known greenhouse gas (1). For example, autotrophic bacteria can use CO_2_ as a carbon source to synthesize the biodegradable plastic poly(3-hydroxybutyrate) (PHB) (2). PHB is a promising sustainable alternative to petroleum-based plastics due to its biodegradability (3). Furthermore, to manufacture value-added products using these bacteria, disrupting PHB synthesis genes to suppress its accumulation and introducing exogenous genes to divert intermediates toward useful materials represents a promising strategy (4). For example, Schoenmakers et al. (2025) disrupted genes, including *phaB*, which encodes acetoacetyl-CoA reductase in the PHB metabolic pathway, and introduced exogenous genes for isopropanol synthesis in a hydrogen-oxidizing bacterium, *Cupriavidus necator*, thereby suppressing PHB accumulation and enabling isopropanol production from CO_2_ as the sole carbon source (5).

*Hydrogenophilus thermoluteolus* TH-1 is a facultative chemoautotrophic thermophilic hydrogen-oxidizing bacterium (6). Its optimal pH and temperature are 7.0 and 52°C, respectively. The doubling time of this strain under autotrophic conditions is approximately 1 hour (μ = 0.68), placing it among the fastest-growing autotrophic bacteria. Furthermore, TH-1 can accumulate PHB under nitrogen-deficient conditions regardless of autotrophic or heterotrophic growth (7). Therefore, by combining high culture temperature, high CO_2_ fixation capacity, and a bioengineered PHB synthesis pathway, strain TH-1 can be exploited to produce useful biomaterials from CO_2_ with minimal contamination.

Three key enzymes drive the main PHB synthesis pathway: β-ketoacyl-CoA thiolase (PhaA), which condenses two acetyl-CoA molecules to acetoacetyl-CoA; acetoacetyl-CoA reductase (PhaB), which reduces acetoacetyl-CoA to (*R*)-3-hydroxybutyryl-CoA; and PHB synthase (PhaC), which polymerizes (*R*)-3-hydroxybutyryl-CoA to PHB (8). In addition to this primary pathway, PHB synthesis involves β-oxidation (9). Crotonyl-CoA, a β-oxidation intermediate, is converted by *R*-specific 3-hydroxybutyryl-CoA hydratase to (*R*)-3-hydroxybutyryl-CoA, which is then polymerized into PHB by PhaC (10). Because acetoacetyl-CoA is a key intermediate that can be diverted to useful materials such as isopropanol, *phaB* genes are frequently disrupted to suppress PHB synthesis (5). Genome analysis of strain TH-1 revealed that PhaA, PhaB, and PhaC are involved in its PHB synthesis pathway (11). Several genes encoding enzymes that catalyze the same step of the synthetic pathway have been identified, including *phaA1-3, phaB1-2*, and *phaC1-3* (7).

In this study, we disrupted the acetoacetyl-CoA reductase genes, *phaB1* and *phaB2*, as a foundational step toward redirecting carbon flux from CO_2_ toward the production of other value-added biomaterials. As expected, the disruption of these genes significantly reduced PHB accumulation; however, the resulting mutant still retained the ability to synthesize PHB at a detectable level. This phenotype further encouraged us to investigate the metabolic flexibility of strain TH-1. Here, we demonstrate the existence of an alternative, PhaB-independent PHB synthesis pathway and discuss the involvement of (*R*)-specific fatty acid β-oxidation intermediates in this bypass mechanism under both autotrophic and heterotrophic conditions.

## MATERIALS AND METHODS

### Bacterial strains and growth media

*H. thermoluteolus* TH-1 (DSM 6765) wild-type and its Δ*phaB1B2* mutant were used in this study. A nitrogen-containing inorganic medium was prepared by adding (NH_4_)_2_SO_4_ (3 g), KH_2_PO_4_ (1 g), K_2_HPO_4_ (2 g), MgSO_4_•7H_2_O (0.5 g), NaCl (0.25 g), CaCl_2_ (0.03 g), FeSO_4_•7H_2_O (0.014 g), and 50 μL trace element solution to 1 L of distilled water, followed by sterilization. The trace element solution contained MoO_3_ (40 mg/L), ZnSO_4_•7H_2_O (280 mg/L), CuSO_4_•5H_2_O (20 mg/L), H_3_BO_3_ (40 mg/L), MnSO_4_•5H_2_O (40 mg/L), and CoCl_2_•6H_2_O (40 mg/L). A medium lacking (NH_4_)_2_SO_4_ was used as a nitrogen-deficient inorganic medium to promote PHB accumulation in strain TH-1. The cell growth was monitored by measuring the optical density at 600 nm (OD_600_) using a spectrophotometer (Beckman Coulter, USA).

Bacteria were grown under autotrophic conditions. First, the glycerol stock was inoculated into 5 mL of nitrogen-containing inorganic medium in a test tube, which was then sealed. The headspace was replaced with a gas mixture of H_2_:O_2_:CO_2_ (75:10:15) and incubated overnight at 50°C and 300 rpm. Next, the resulting culture was inoculated into a 50-mL fresh nitrogen-containing inorganic medium in a 500-mL medium bottle. The bottle was sealed, and the gas was replaced as described above. The culture was incubated overnight at 50°C and 300 rpm until it reached the late logarithmic or early stationary phase.

PHB accumulation under autotrophic conditions was quantified as follows: The 50-mL culture obtained above was centrifuged to harvest the cell pellet, which was washed once with a nitrogen-deficient inorganic medium. Then, one fifth of the pellet was inoculated into 10 mL of fresh nitrogen-deficient inorganic medium in a 100-mL vial, which was sealed. After replacing the gas mixture, the culture was incubated for 24 hours at 50°C and 300 rpm. The resulting culture was centrifuged to harvest the cell pellet, which was divided into equal portions. The portions were heated for 20 minutes at 100°C using a heat block to obtain the dried biomass, which was analyzed by high-performance liquid chromatography (HPLC) and gas chromatography (GC).

To quantify PHB accumulation under heterotrophic conditions, cells grown under autotrophic conditions were harvested by centrifugation, washed once with a nitrogen-deficient inorganic medium, and then inoculated into 5 mL aliquots of nitrogen-deficient medium containing one of the following carbon sources—20 mM sodium acetate, 40 mM disodium DL-malate, 20 mM sodium butyrate, and 2 mM sodium oleate. After incubation for 4 hours at 50°C and 300 rpm, the resulting culture was harvested by centrifugation, dried, and analyzed by HPLC.

### The Δ*phaB1B2* strain construction

To create the Δ*phaB1B2* mutant strain, a plasmid was synthesized with disrupted *phaB1* and *phaB2*, which are located tandemly on the chromosome, by GenScript (USA). The upstream region of *phaB1*, a kanamycin resistance gene (*nptII*) codon-optimized for strain TH-1, and the downstream region of *phaB2* (Supplementary Text) were cloned between the EcoRI and BamHI sites of pUC57. The plasmid was linearized by EcoRI digestion before electroporation.

To prepare the competent TH-1 strain, cells were cultivated under autotrophic conditions and harvested at the middle logarithmic phase. The harvested cells were washed with deionized water and 10% glycerol. Then, linearized DNA was added to the cells, which were electroporated using a Gene Pulser (Bio-Rad, USA) under exponential decay conditions (25 µF capacitance, 100 Ω resistance, 2,900 V applied voltage). Deletion mutants obtained via double-crossover homologous recombination were selected on agar plates containing an inorganic medium supplemented with 50 μg/mL kanamycin under a gas phase of H_2_:O_2_:CO_2_ (75:10:15). The gene disruption was confirmed by PCR and DNA sequencing.

### Transmission electron microscope (TEM) analysis

The Δ*phaB1B2* strain was inoculated into 100 mL nitrogen-containing inorganic medium in a 1 L medium bottle and incubated overnight at 50°C and 130 rpm until the culture reached the late logarithmic or early stationary phase. The resulting culture was harvested by centrifugation and washed once with a nitrogen-deficient inorganic medium. Then, the cells were inoculated into 15 mL aliquots of nitrogen-deficient medium containing different carbon sources, including 10 mM disodium DL-malate, 10 mM butyric acid, or 2.2 mM sodium oleate, in 250 mL medium bottles and incubated for 5 hours at 50°C and 140 rpm. The cells were fixed using fixing solution A (4 % paraformaldehyde and 4 % glutaraldehyde in 0.1 M cacodylate buffer at pH 7.4) and fixing solution B (2 % glutaraldehyde in 0.1 M cacodylate buffer at pH 7.4), following the protocol provided by Tokai Electron Microscopy, Inc. TEM analysis was conducted by Tokai Electron Microscopy, Inc.

### HPLC and GC analyses

The HPLC analysis was performed as described previously (7, 12). PHB content was determined based on the conversion to crotonic acid through acid-catalyzed elimination during chemical depolymerization. After adding concentrated sulfuric acid to the dried cells, the mixture was heated at 100°C for 1 h using a heat block. The resulting crotonic acid was measured using an HPLC system equipped with a TSKgel OApak-A ion-exclusion column (7.8 mm × 30 cm; Tosoh, Tokyo, Japan). Using 0.75 mM H_2_SO_4_ as the mobile phase at a flow rate of 0.6 mL/min, crotonic acid was detected by monitoring absorbance at 210 nm. Its concentration was expressed as mol/L culture per OD_600_ (mM/OD_600_).

GC analysis was performed using a previously described method with some modifications (13). Methanolysis was performed for GC derivatization. Dried cells were added to a 1 mL mixture of chloroform and methanol (1:1, v/v) in screw-cap test tubes. The methanol used for the mixture contained 3% (v/v) concentrated sulfuric acid, and the final mixture contained 0.25 mg/mL benzoic acid as an internal standard. The test tubes were heated at 100°C for 6 hours, and PHB was converted to 3-hydroxybutyrate methyl ester (3HB ME). After cooling the mixtures to room temperature, 0.25 mL of a solution containing 1 M NaCl and 1.32 M NaOH was added to the reaction solutions, mixed for 10 minutes at 250 rpm, and centrifuged for 3 minutes at 15,000 rpm. The organic phase was collected and analyzed using the DB-23 GC column (length: 30 m; diameter: 0.250 mm; film: 0.25 μm) (Shimadzu, Japan) with a flame ionized detector. The start temperature was 100°C and the hold time was 0 minutes. The temperature was ramped at 20°C/min for 5 minutes, and the final temperature of 200°C was held for 1 minute. 3HB ME was used as an external standard. The concentration of 3HB ME was expressed as mol/L of culture per OD_600_ (mM/OD_600_).

## RESULTS

### Analysis of PHB accumulation by Δ*phaB1B2*

The *phaB1B2* genes were disrupted by homologous recombination, and the gene deletion was confirmed by PCR and DNA sequencing (data not shown). This deletion did not affect cell growth under normal conditions (data not shown).

The wild-type and Δ*phaB1B2* strains were cultured in a nitrogen-deficient inorganic medium under autotrophic conditions. HPLC analysis was employed to quantify PHB by converting it into crotonic acid via acid hydrolysis. The analysis revealed that the crotonic acid concentration was 1.25±0.04 mM/OD_600_ for the wild-type, while the concentration was 0.32±0.02 mM/OD_600_ for the Δ*phaB1B2* strain, which was approximately 25 % of that in the wild-type strain (Fig. 1A). This result demonstrated that disruption of *phaB1* and *phaB2* markedly reduced PHB accumulation in strain TH-1, but suggested that PHB or related metabolites convertible to crotonic acid are still synthesized in the mutant strain.

**FIG. 1.**
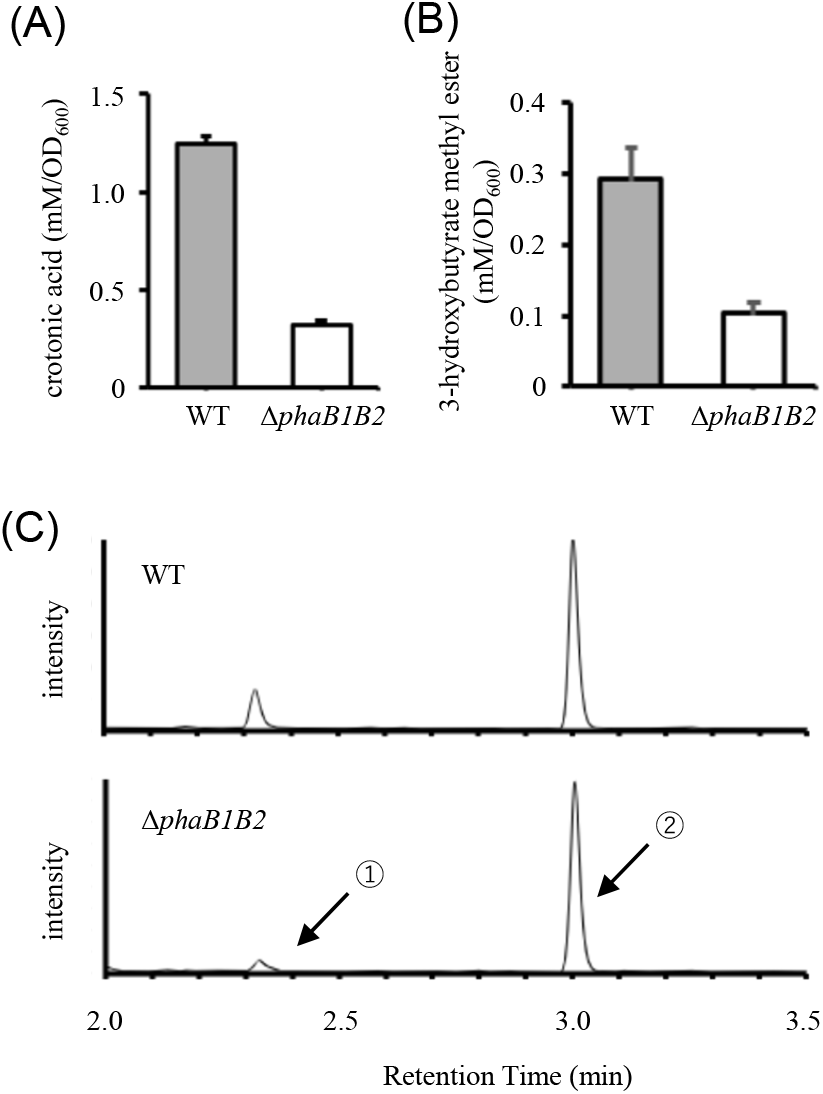
(A) Crotonic acid and (B) 3-hydroxybutyrate methyl ester concentrations in the wild-type and ΔphaB1B2 strains under autotrophic conditions quantified by HPLC and GC, respectively (n = 3; error bars represent standard deviation). (C) Representative chromatograms corresponding to (B). Peaks 1 and 2 indicate 3-hydroxybutyrate methyl ester, and benzoic acid methyl ester (internal standard), respectively.

To determine if the crotonic acid measured by HPLC in the Δ*phaB1B2* strain was derived from PHB or a monomeric soluble intracellular compound, the cells of the wild-type and Δ*phaB1B2* strain cultured in a nitrogen-deficient inorganic medium under autotrophic conditions were disrupted by sonication. The lysate was then separated into a cell-free extract and the insoluble fraction. Subsequently, HPLC analysis was performed on both fractions to measure the crotonic acid concentration. As a result, crotonic acid was detected only in the insoluble fraction (Supplementary Fig. 1). Therefore, the crotonic acid detected under the nitrogen-starvation conditions is most likely derived from PHB.

To confirm that the crotonic acid measured by HPLC was derived from PHB, methanolysis was conducted for the wild-type and Δ*phaB1B2* strains, and the resulting 3HB ME was measured by GC. Peaks corresponding to 3HB ME were observed in the GC chromatograms of both strains (Fig. 1C), and their identities were confirmed by GC/MS (data not shown). The 3HB ME concentration was 0.29±0.04 mM/OD_600_ in the wild-type strain, while in the Δ*phaB1B2* strain, it was 35 % of this value (0.10±0.02 mM/OD_600_) (Fig. 1B). These results clearly indicated that the Δ*phaB1B2* strain could produce PHB.

### PHB accumulation in the wild-type and Δ*phaB1B2* strains using acetate, malate, butyrate, or oleate as carbon sources

To explore the contribution of β-oxidation to PHB synthesis, we investigated the effect of the addition of fatty acids on PHB accumulation. The fatty acids butyrate and oleate were used as carbon sources for cell culture under nitrogen-starvation conditions. Acetate and malate were selected as carbon sources in the control group, as they are catabolized to acetyl-CoA, thereby promoting PHB synthesis via the PhaB pathway. When grown on acetate and malate, the crotonic acid concentrations in the Δ*phaB1B2* strain were only 17% and 26% of those in the wild-type strain, respectively (Fig. 2). In contrast, when butyrate and oleate were supplied, the concentrations in the Δ*phaB1B2* strain reached 72% and 80% of those in the wild-type strain, respectively. These results indicate that PHB accumulation was significantly restored in the Δ*phaB1B2* strain by the addition of fatty acids.

**FIG. 2.**
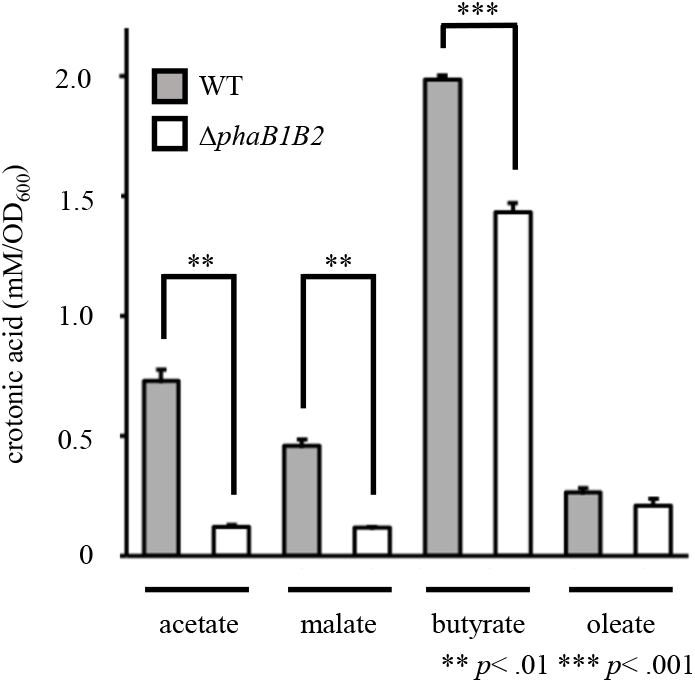
Concentrations of crotonic acid in the wild-type and Δ*phaB1B2* strains supplied with acetate, malate, butyrate, or oleate (n = 3; error bars represent SD). \*\**p* <.01, \*\*\**p* <.001

TEM analysis revealed significant differences in PHB granule formation depending on the carbon source provided to the Δ*phaB1B2* strain (Fig. 3). When cultivated with butyrate (Fig. 3C, D) or oleate (Fig. 3E, F), the cells exhibited prominent formation of intracellular PHB granules. In contrast, when supplied with malate, only a few small granules were observed in a minority of the cells (Fig. 3A, B). These morphological observations are consistent with the quantitative results, which showed that PHB accumulation in the Δ*phaB1B2* strain was markedly higher when fatty acids were supplied compared to when malate was used (Fig. 2).

**FIG. 3.**
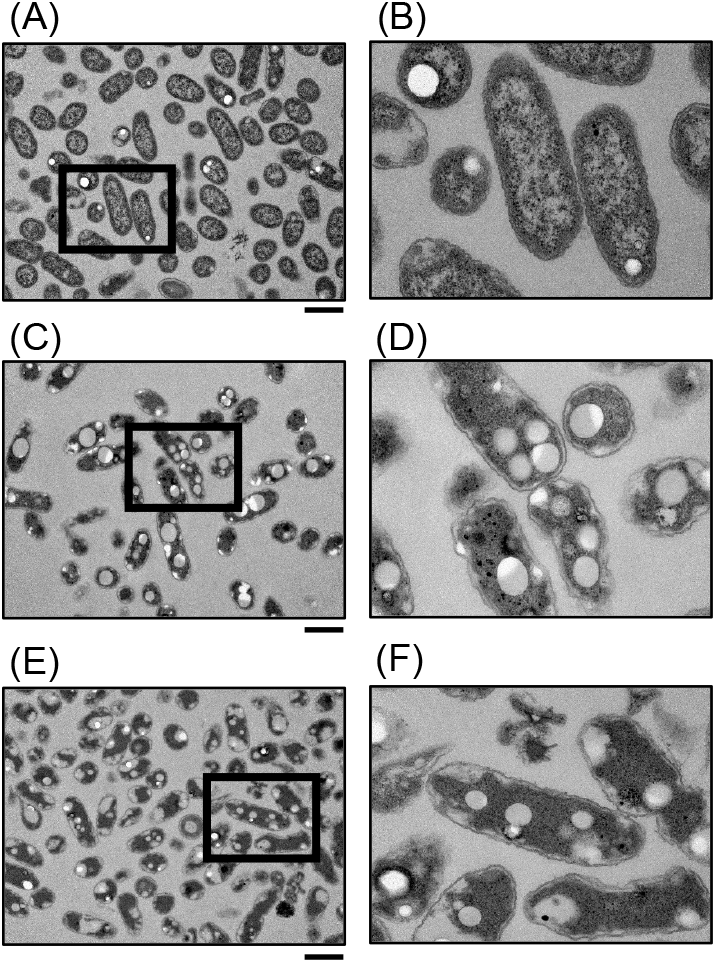
TEM images of the Δ*phaB1B2* strain supplied with malate (A, B), butyric acid (C, D), or oleate (E, F). Panels B, D, and F are enlarged views of the boxed regions in A, C, and E, respectively. Scale bars: 1 μm.

## DISCUSSION

In this study, PHB accumulation in the Δ*phaB1B2* strain was significantly reduced under autotrophic conditions, as evidenced by the HPLC, GC, and TEM analyses. This observation supports the role of PhaB1 and PhaB2 as the primary enzymes involved in PHB synthesis in this strain. However, the Δ*phaB1B2* strain still accumulated a considerable amount of PHB, suggesting the presence of a PhaB-independent PHB synthesis pathway in this strain.

Higher PHB accumulation was observed in the Δ*phaB1B2* strain when butyrate or oleate was used as a carbon source than when acetate or malate. These results suggest that the residual PHB accumulation in the Δ*phaB1B2* strain is specifically linked to the metabolism of fatty acids. We propose a PhaB-independent PHB synthesis pathway in strain TH-1 (Fig. 4). While the contribution of β-oxidation to PHB accumulation has been documented in other bacteria (14-16), this study represents the first demonstration of such a bypass mechanism in the fast-growing hydrogen-oxidizing bacterium *H. thermoluteolus*. When butyrate and oleate are metabolized via β-oxidation, intermediates such as crotonyl-CoA and (*S*)-3-hydroxybutyryl-CoA are produced. These intermediates can be converted to (*R*)-3-hydroxybutyryl-CoA, which serves as the substrate for PhaC. Therefore, we hypothesize that strain TH-1 possesses the genes that facilitate to the production of (*R*)-3-hydroxybutyryl-CoA from these intermediates.

**FIG. 4.**
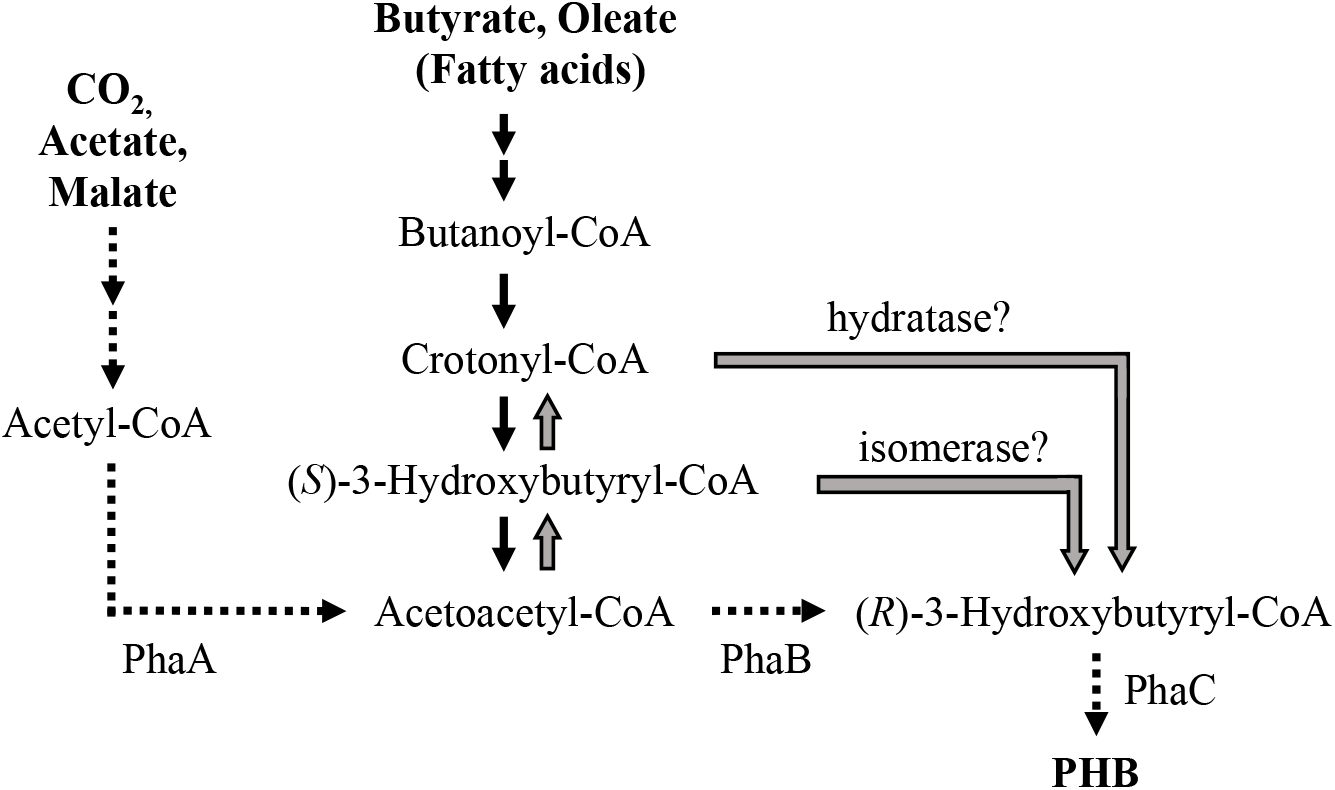
The proposed PHB synthesis pathway in strain TH-1. Dashed arrows indicate the PhaB-dependent pathway. Black solid arrows indicate the β-oxidation pathway. Gray solid arrows indicate PhaB-independent pathway via reverse β-oxidation.

Furthermore, the fact that Δ*phaB1B2* strain accumulates PHB using CO_2_, acetate, or malate suggests that PHB was produced via the reverse flow of the β-oxidation in strain TH-1. For the synthesis of PHB from these carbon sources, one of the most plausible pathways is the conversion to acetoacetyl-CoA and then to crotonyl-CoA or (S)-3-hydroxybutyryl-CoA through reverse β-oxidation. This indicates that strain TH-1 may possess a native reverse β-oxidation pathway, as observed in other bacteria (17).

In conclusion, this study sought to suppress PHB accumulation as a critical step toward establishing strain TH-1 as a versatile microbial platform. While the disruption of *phaB1B2* significantly reduced PHB levels, the unexpected persistence of synthesis revealed a robust metabolic flexibility. These findings indicate that complete carbon flux redirection will require the inactivation of not only the primary PhaABC pathway but also these secondary routes. Future engineering of the β-oxidation-related genes identified here will be essential to fully harness the potential of strain TH-1 for efficient CO_2_-based biomanufacturing.

## Acknowledgments

This work was supported in part by the commissioned research project “Transformation of the Aquaculture Industry into a Growth Industry,” funded by the Fisheries Agency, Ministry of Agriculture, Forestry and Fisheries (MAFF), Japan.

