## Supplemental Information for "PhaB-independent poly(3-hydroxybutyrate) production in the thermophilic hydrogen-oxidizing bacterium *Hydrogenophilus thermoluteolus*"

**Abstract:**

*Hydrogenophilus thermoluteolus* TH-1 is a thermophilic hydrogen-oxidizing bacterium capable of producing poly(3-hydroxybutyrate) (PHB) from $CO_2$. To redirect carbon flux for producing other useful biomaterials, we disrupted the acetoacetyl-CoA reductase genes (*phaB1* and *phaB2*), which are central to the primary PHB synthesis pathway. Unexpectedly, the resulting Δ*phaB1B2* mutant still accumulated PHB under autotrophic conditions, reaching approximately 25-35 % of the wild-type level. Furthermore, PHB accumulation in the mutant was significantly restored when fatty acids (butyrate and oleate) were used as carbon sources, whereas acetate and malate resulted in reduced accumulation. These results suggest the existence of a PhaB-independent PHB synthesis pathway. We propose that intermediates from the β-oxidation of fatty acids are converted to (*R*)-3-hydroxybutyryl-CoA, bypassing the disrupted PhaB enzymes. Additionally, the basal PHB production from non-fatty acid sources implies the involvement of a reverse β-oxidation pathway. This study highlights the metabolic versatility of strain TH-1 for future metabolic engineering.

### SUPPLEMENTARY DATA

#### Legend to Supplementary Figure

Supplementary FIG. 1. Crotonic acid concentrations in whole cells, insoluble fractions, and cell-free extracts in the wild-type and Δ*phaB1B2* strains cultured under autotrophic conditions with or without nitrogen sources (ammonium sulfate), as quantified by HPLC.

Supplementary Fig. 1

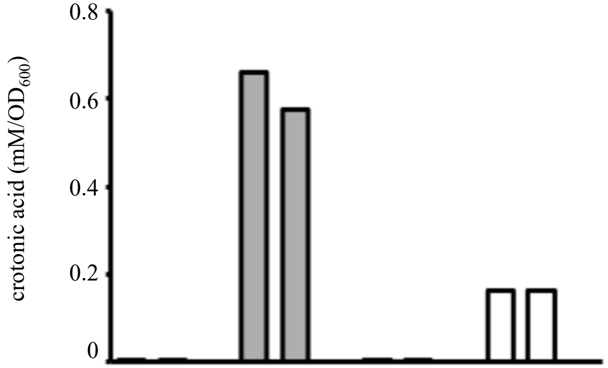


| strain | WT | | | | | | *ΔphaB1B2* | | | | | |
|---|---|---|---|---|---|---|---|---|---|---|---|---|
| nitrogen | + | | | - | | | + | | | - | | |
| whole cell | + | - | - | + | - | - | + | - | - | + | - | - |
| Insoluble fraction | - | + | - | - | + | - | - | + | - | - | + | - |
| cell-free extract | - | - | + | - | - | + | - | - | + | - | - | + |

>Supplementary Text. The nucleotide sequence of the insertion fragment used to disrupt phaB1 and phaB2.

GAATTCCGGCTGGGACGACTACCTGGCGTTGGGGCCGCTCACAGCGCTCGAAGTCGTGCGGGAAATC
ACCGGTTGTCCCGATCCCAACGTGTTGGGATTCTGCGTCGGCGGAACGATCCTGGCGTCGGCGCTCG
CCGTTGCGGTTGCCCGTGGCGATCCGCCCGTGCATAGTCTGACCCTGATGACGACACTCTTGGATTT
TGCCGAAAGCGGGGAACTGGGGTGCCTGGTCGATGAGGCGAGCGTTGCGGCGCGGGAGGCGACGATC
GGCCAAGGTGGCTTGATGTTGGGGTCGGAACTGGCGCAAGTCTTTTCGATGTTACGCGCGAACGACC
TCATTTGGAATTACGTCGTCGGTAACTATTTGAAGGGGGAAAAACCCAAGCCCTTCGATCTGCTCTA
TTGGAACAGCGACAGCACCAATCTGCCCGGGCCGTTCGTCACCTACTACTTGCGCCATATGTATCTG
GAAAACGAACTCCGCCTGCCCAACAAACTGAAAATGCTCGGGTACCCCGTCAATCTCGGTGCCATCA
CCGCGCCCGCCTACGTAATGGCGGCGCGGGAAGACCACATCGTTCCGTGGCGCGGGGCATACCATTC
GATGCAGCTTTTGGGGGGTGAAAAACGGTTCGTGTTGGGCGCAAGCGGTCATATCGCGGGTGCGATC
AATCCCGCGAAGAAGAATCGGCGCAGTTATTGGGTCAATCCCGAATTGCCGCAGGAGCCGGACGAAT
GGTTTGCCGCGGCGCGCGAATGCCCCGGGAGTTGGTGGAACGATTGGGCCGATTGGCTGAAGGAGCA
CAAAGGGAAAACGGTGCCTGCGCGTGAGTCGCTCGGAAACGACACATACCCACCGATCGAACCCGCT
CCGGGACGCTACGTCAAAGAACCGGCGCTGCCGCTCGTGCGAGGATCGCGTGAAGCCGCGGCCGTGT
GAGTAGCGGTGGAATCGAAAAAGGGAGGGTCAAATGGTTTTCAACACCACAAGGAGGGAGAGCTCAT
GATCGAACAAGACGGTCTCCACGCGGGGTCGCCGGCGGCGTGGGTCGAACGCCTCTTCGGCTACGAC
TGGGCGCAGCAAACCATCGGGTGCTCGGACGCGGCGGTGTTCCGCCTCAGCGCGCAGGGTCGCCCGG
TCCTCTTCGTGAAAACCGACCTCTCGGGGGCGCTCAACGAACTCCAAGACGAAGCGGCGCGCCTCTC
GTGGCTCGCGACCACCGGCGTCCCGTGCGCGGCGGTGCTCGACGTGGTGACCGAAGCGGGTCGCGAC
TGGCTCCTCCTCGGCGAAGTCCCGGGTCAAGACCTGCTCTCGTCGCACCTCGCGCCGGCGGAAAAAG
TGAGCATCATGGCGGACGCGATGCGCCGCCTCCACACCCTCGACCCGGCGACCTGCCCGTTCGACCA
CCAAGCGAAACACCGCATCGAACGCGCGCGCACCCGCATGGAAGCGGGCCTCGTCGACCAGGACGAC
CTCGACGAAGAACACCAAGGTCTCGCGCCGGCGGAACTCTTCGCGCGCCTCAAAGCGCGCATGCCGG
ACGGGGAAGACCTCGTGGTGACCCACGGGGACGCGTGCCTCCCGAACATCATGGTCGAAAACGGCCG
CTTCTCGGGGTTCATCGACTGCGGCCGCCTCGGGGTGGCGGACCGCTACCAGGACATCGCGCTCGCG
ACCCGCGACATCGCGGAAGAACTCGGCGGGGAATGGGCGGACCGCTTCCTCGTGCTCTACGGTATCG
CGGCGCCGGACAGCCAACGCATCGCGTTCTACCGCCTCCTCGACGAGTTCTTCTGAGCCACTCGTTT
GACGAAACGAGACTTTTTGGCGGCTGGGCTGCGAACGGCGGTATAATTGCCCGATTGCGGCCGAGCC
GCGTGACCATGAGAGTGTTCCATGAACGCGGGAAGCGAACGCGTCATCAAGAAATATCCCAACCGGC
GGCTCTACGATACCGCAACCAGCACCTATATCACGCTGGCGGAAATCAAAGAGATGATCCTTCATCA
CGAGCCGGTGCGTGTGATCGACGCCAAAACGCAAGAAGATCTGACGCGCAGCGTTTTGCTGCAGATT
TTGATGGAAGAAGAGGTGGGGGGAGAGCCCATCTTTTCCACCGAAATGCTCGCCAGTATCATTCGTT
TCTACGGCCAGGCGTCGCAGACGCTTTTTGGTCAATACCTGGAAAACAGCCTGAAAACGTTCCTGGA
ACTGCAAAGCCGCATGCAAGAGCGCTTTCGCGCAATCTACGGTGACAACGCGGTGATGGGGCCAGAG

ATCCTCGCCCAGTTTCTTTCACTGCAACCGAAAGCGATGCAGTCGATGCTTGATGCCTACCTCGAGC
AAACCCAAAACTTATGGCATCAAATGAACCAACAGATGCAGCGCCAAGCGCTGAGCTGGTTCGAAGC
TTTCTATCCCAAATCCAATCCCACCTCGAAAAACCCGCAATCGCAAGATGAATCGAAGTGACCCCAC
GCCGCGCGTCGGCATGGTGAGTTTGGGGTGCCCCAAAGCCACCGTCGACACCGAACGCATTTTGACG
CGACTTCGTGCCGAAGGGTATGAAATCACGCCCGAATACGGCGAAGCCGATCTCGTCATCGTCAATA
CCTGTGGCTTCATCGAAGAAGCCGTGGCAGAGTCGCTCGACGCGATCGGCGAAGCGATCCAGGAAAA
CGGTAAGGTGATCGTCACAGGCTGTTTGGGGGCGCGCCGTGACGTGATCCTTGCAGCCCACCCCAAG
GTGCTTGCGGTGACAGGGCCGCATGCCACCGACGAAGTGATGGCGCTCGGATCC
